# Is proteolytic cleavage essential for the activation of *Hydra* pore-forming toxin, HALT-4?

**DOI:** 10.1101/2022.08.13.503830

**Authors:** Yap Wei Yuen, Hwang Jung Shan

**Affiliations:** Department of Biological Sciences, School of Medical and Life Sciences, Sunway University, Selangor Darul Ehsan, Malaysia; Department of Medical Sciences, School of Medical and Life Sciences, Sunway University, Selangor Darul Ehsan, Malaysia

**Keywords:** *Hydra* actinoporin-like toxin, pore-forming toxin, dibasic cleavage site, regulated secretory pathway, cytolytic activity

## Abstract

The mature form of *Hydra* actinoporin-like toxin 4 (mHALT-4) differs from other actinoporins primarily by bearing approximately 115 additional residues at the N-terminus. Five dibasic residues were found in this extended region and we assume that, when cleaved, each truncated HALT-4 (tKK1, tKK2, tRK3, tKK4 and tKK5) could exhibit an enhanced cytolytic activity. However, our results showed that mHALT-4, tKK1 and tKK2 possessed similar cytolytic activity against HeLa cells, whereas tRK3, tKK4 and tKK5 failed to kill HeLa cells. Therefore, the cleavage of KK1 or KK2 sites did not enhance the cytolytic activity of tKK1 and tKK2 but might facilitate the sorting of tKK1 and tKK2 to the regulated secretory pathway and eventually deposit them in the nematocyst. In contrast, RK3, KK4 and KK5 were unlikely to serve as the proteolytic cleavage sites since the amino acids between KK2 and RK3 are also crucial for the pore formation.

**Highlights:**

- Five dibasic cleavage sites are identified at the N-terminal region of HALT-4 but they are unlikely to have a role in the enhancement of HALT-4 cytotoxicity.
- The first two dibasic sites, KK1 and KK2, may be involved in sorting HALT-4 to the regulated secretory pathway where HALT-4 can be transported to the nematocyst.

---

*Hydra* actinoporin-like toxin (HALT) is a family of seven α-pore-forming toxins, namely HALT-1 to HALT-7 (Glasser et al., 2014; Yap et al., 2019). As their name suggests, their secondary and tertiary structures highly resemble that of actinoporins of sea anemones, especially HALT-1, 5, 6 and 7. HALT-3 and HALT-4 deviate from actinoporin by having a shorter and an extended N-terminal peptide, respectively (Yap et al., 2019). Actinoporin is stored as reservoir in nematocyst, which is a capsular organelle with a long coiled thread, and is believed to play an offensive role in hunting preys. Similarly, HALTs-1, 4 and 7 were also detected in the nematocyst-containing cells, however, HALT-1 as well as HALT-2 and 6 were also detected in other cell-types (Yap et al., 2019).

Except HALT-3, all HALTs contain a signal peptide at the *N-*terminal end. Signal peptide is meant to be cleaved when HALT proteins are processed in the rough endoplasmic reticulum before HALT becomes a mature protein. According to Glasser et al. (2014), HALT-4 and HALT-2 contain a putative propart because of the extended N-terminal peptide, while other HALTs contain 0-3 residues between the signal peptide and the mature polypeptide. As compared to mature HALT-1, the putative propart of mature HALT-4 (mHALT-4) has 115 additional amino acids and this region contains 5 dibasic sites (KK or KR) (Rholam et.al 1986; Rholam et.al 1995). Therefore, HALT-4 is unique among other HALTs as HALT-4 contains a propart-like sequence that could be cleaved for protein maturation. The 5 potential cleavage sites at the N-terminal region of HALT-4 are shown in Figure 1A. The dibasic cleavage sites is not found in HALT-2. Many cnidarian toxins including equinatoxin II contain a signal peptide and followed by 9-17 residues of propart at the N-terminus. Anderluh et al. (2000) have revealed that a number of genes encoding nematocyst-associated proteins and toxins possess highly conserved signal peptide and propart ending with KR dibasic residues. For example, residues 1-19 of equinatoxin II is the signal peptide which directs the precursor polypeptide to ER and the following residues 20-35 is the propart with KR as the last two residues (Belmonte et al., 1994; Anderluh et al., 1996). Moreover, the homolog of prohormone convertase 3 (PC3) of vertebrates was also found in *Hydra. Hydra* PC3-like protein is 55.4% and 56.7% identical in the catalytic domain to mouse PC3 and human furin, respectively (Chan et al., 1992). This finding suggested that the *Hydra* PC3-like protein may process the precursor form of HALT-4 by targeting and cleaving one or more dibasic site(s) at the N-terminus of HALT-4. The mHALT-4, when expressed in a recombinant form, was capable of killing human cells but its activity was seven-fold weaker than the recombinant EqtII (Glasser et al., 2014; Yap et al., 2019). Therefore, our work considered the possibility that HALT-4 can be enzymatically cleaved at one of the dibasic sites, and the cleaved HALT-4 would have a significantly enhanced cytotoxicity.

**Figure 1.**
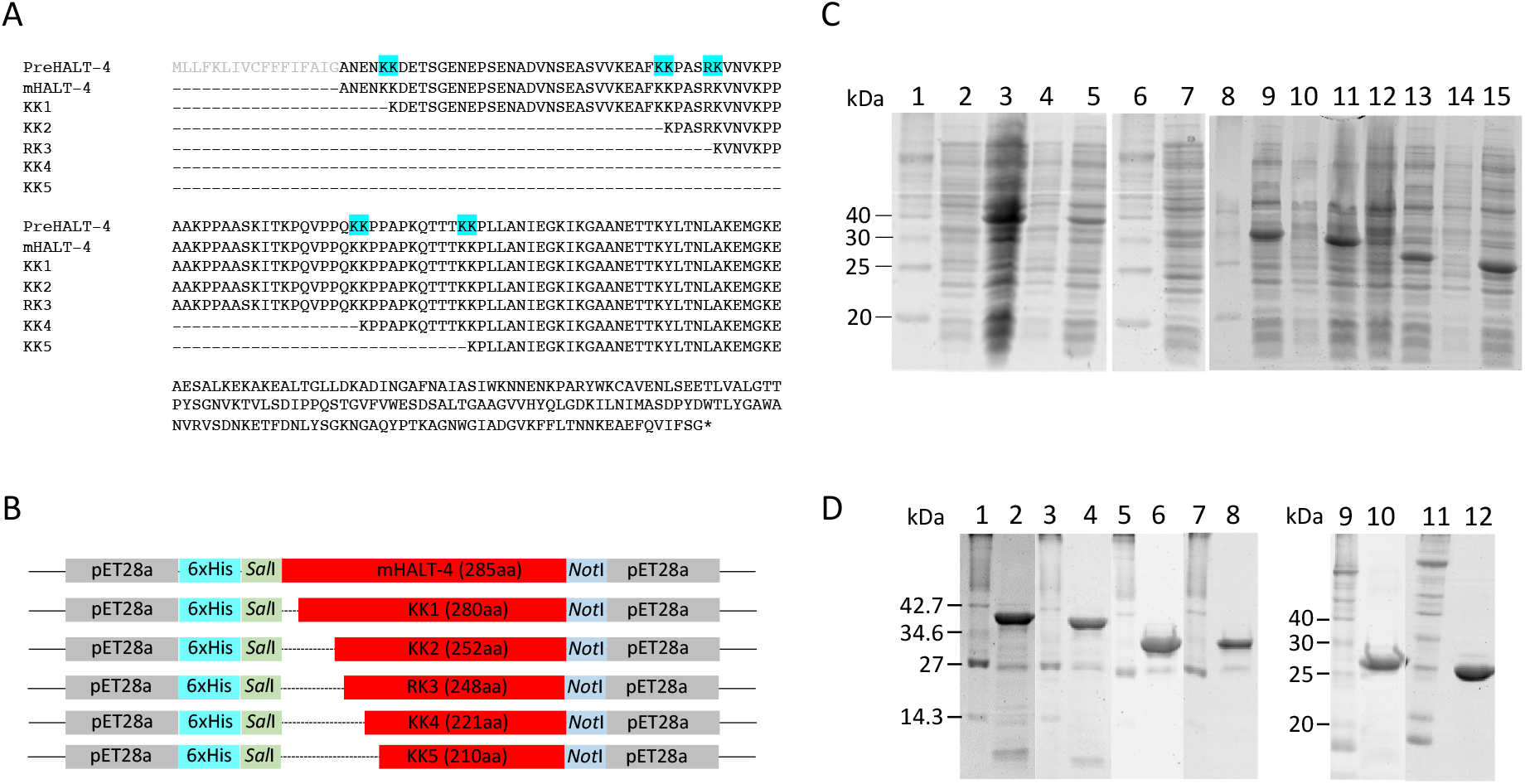
(A) N-terminal amino acid sequence alignment of HALT-4 precursor (PreHALT-4), mature HALT4 (mHALT-4) and various truncated HALT-4 (KK1, KK2, RK3, KK4 and KK5). Five dibasic sites are highlighted in light blue. (B) Schematic drawing of the orientation of recombinant mHALT-4 and truncated HALT-4 in pET28a. Dotted lines indicate the removal of amino acids in HALT-4. (C) 12% SDS-PAGE of the expression of mHALT-4 and truncated HALT-4 in the presence of 3% ethanol. Lanes 1, 6 and 8, protein markers; lane 2, mHALT-4 without IPTG; lane 3, mHALT-4 with IPTG (35.7 kDa); lane 4, KK1 without IPTG; lane 5, KK1 with IPTG (34.75kDa); lane 7, KK2 without IPTG; lane 9, KK2 with IPTG (31.76 kDa); lane 10, RK3 without IPTG; lane 11, RK3 with IPTG (31.22 kDa); lane 12, KK4 without IPTG; lane 13, KK4 with IPTG (28.52 kDa); lane 14, KK5 without IPTG; lane 15, KK5 with IPTG (27.34 kDa). (D) 12% SDS-PAGE of mHALT-4 and various truncated HALT-4 after Nickel affinity purification and dialysis. Lanes 1, 3, 5, 7, 9 and 11, protein markers; lane 2, mHALT-4; lane 4, KK1, lane 6, KK2; lane 8, RK3; lane 10, KK4 and lane 12, KK5.

Assuming each of the five dibasic sites can be possibly cleaved to generate mature HALT-4, we used mHALT-4 as a template to construct five truncated HALTs-4, namely tKK1, tKK2, tRK3, tKK4 and tKK5 (Figure 1B). Each truncated HALT-4 having the propart removed at a specific dibasic site was cloned into pET28a and transformed into DH5α before sequencing. After the confirmation of truncated HALT-4 sequences, together with the mHALT-4, they were transformed into Rosetta Gami 2 (DE3) *E. coli*. Recombinant mHALT-4, tKK1, tKK2, tRK3, tKK4 and tKK5 were then expressed in the presence of 1mM IPTG and 3% ethanol (Figure 1C). In Figure 1C, the result showed that the recombinant mHALT-4, tKK1, tKK2, tRK3, tKK4 and tKK5 could be expressed. The expected molecular weight of mHALT-4, tKK1, tKK2, tRK3, tKK4 and tKK5 were 34.75 kDa, 31.76 kDa, 31.22 kDa, 28.52 kDa, 27.34 kDa and 23.9 kDa, respectively.

Subsequently, these recombinant proteins were purified in denaturing conditions. 8M of urea was included in the cell disruption buffer as well as the binding buffer of Ni-NTA resins. Decreasing concentrations of urea (8M, 6M, 4M, 2M, 1M and 0M) were used to refold protein in Ni-NTA column during the washing steps. Finally, protein was eluted from the column in the presence of high imidazole concentration. Fractions of eluted protein were pooled and dialysed before storing at -80°C. Figure 1D showed the recombinant mHALT-4, tKK1, tKK2, tRK3, tKK4 and tKK5 after the denaturing purification and the dialysis.

To determine which truncated HALT-4 (tKK1, tKK2, tRK3, tKK4 and tKK5) exhibited an enhanced cytolytic activity, all truncated HALTs-4 were subjected for cytotoxicity assays. Together with mHALT-4, tKK1, tKK2, tRK3, tKK4 and tKK5 were separately applied to HeLa cells at various concentrations (5 µg/mL, 10 µg/mL, 15 µg/mL, 20 µg/mL, 25 µg/mL and 30 µg/mL) and examine their cytolytic activity. Since the mHALT-4 exerts the cytolytic activity, we assumed that the proteolytic cleavage at the dibasic site of HALT-4 would be essential for either its enhanced activity in HeLa cells or. When HALTs reduce cell viability by 50%, they were deemed to be cytolytic (CC50). In Figure 2, mHALT-4 (CC50 = ∼21µg/ml), tKK1 (CC50 = ∼23µg/ml) and tKK2 (CC50 = ∼18µg/ml) had exhibited the cytolytic activity towards HeLa cells (Figures 2A and 2B); whereas, tRK3, tKK4 and tKK5 showed no sign of cytolytic activity (Figures 2C, 2D and 2E). The results confirmed that mHALT-4 possessed cytolytic activity as we previously observed (Yap et al., 2019). tKK1 and tKK2 possessed similar cytotoxic level as that of mHALT-4 (Figures 2A and 2B). The polypeptide length of tKK1 and tKK2 are 5 and 33 amino acids shorter than that of mHALT-4, suggesting that this N-terminal short fragment could be negligible in term of its structural and functional roles for HALT-4. Nevertheless, we cannot rule out the possibility that the proteolytic cleavage at KK1 and KK2 sites facilitated the sorting of tKK1 and tKK2 respectively to the nematocyst capsule via the regulated secretory pathway (Anderluh et al., 2000). It has been reported that the dibasic residues act as a sorting signal for targeting secretory protein to the regulated secretory pathway that then releases the secretory protein to its final subcellular location (Brakch et al., 2002; Feliciangeli et al., 2001). Brakch et al. (1994) has reported that a single mutation of the Arg-Lys dibasic could prevent prosomatostatin from entering the regulated secretory pathway. The remaining 77-82 amino acids should be crucial since tRK3, tKK4 and tKK5 did not affect the cell viability in the MTT assays (Figures 2C, 2D and 2E). In fact, the N-terminal region of actinoporins was critical for the initiation of pore formation by inserting the N-terminal domain in the membrane (Kristan et al. 2009). Moreover, the pore formation of sticholysin, a typical actinoporin found in the sea anemone *Stichodactyla helianthus*, was facilitated by the thickness of lipid bilayer (Palacios-Ortega et al. 2017). Based on the findings in the paper, the bilayer thickness of di-18:1 phosphatidylcholine gave the best optimal condition for the penetration of N-terminal α-helix of sticholysins and subsequently the pore formation (Palacios-Ortega et al. 2017). Therefore, HALT-4 having a long N-terminal end could be ideally dedicated to a particular cell type of organism with distinct thickness of lipid bilayer. Removing more than 33 amino acids from the N-terminal end might destabilize the electrostatic and covalent bonds between N-terminal α-helix and the fatty acid chains.

**Figure 2.**
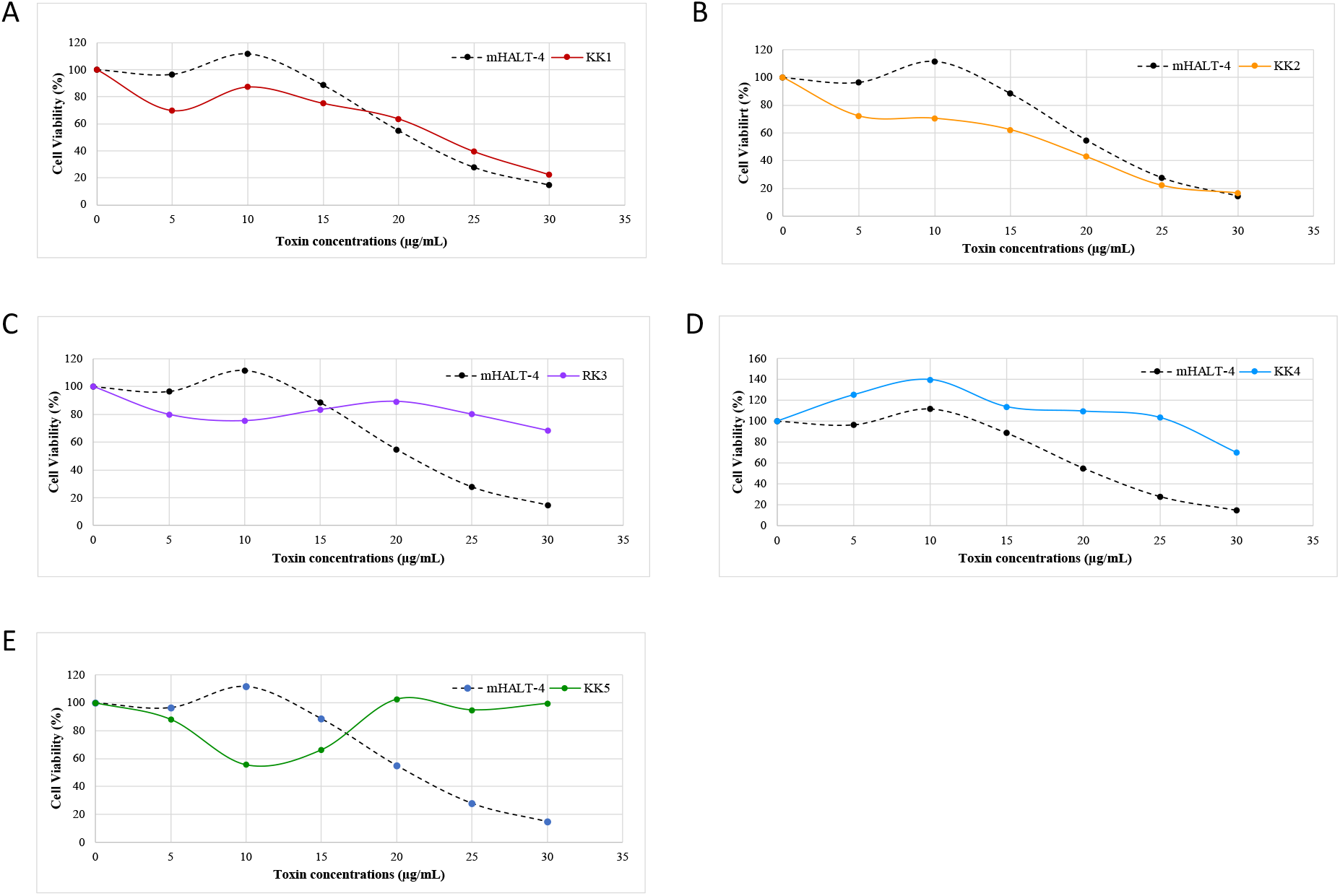
HeLa cells viability after treatment with different concentrations of mHALT-4, and truncated HALTs-4. HeLa cells were incubated with each of recombinant proteins at concentration of 5, 10, 15, 20, 25 and 30 µg/mL at 37°C for 24 hours. Positive control was the cells treated with DMSO and negative control was the cells without treatment. The cell viability was quantified at 570nm/630nm. Each experiment was done in triplicate.

Based on the present findings, we could conclude that RK3, KK4 and KK5 are not the proteolytic cleavage sites and the N-terminal region of HALT-4 is meant to be longer than that of other HALTs and actinoporins. As for KK1 and KK2, the cleavage at these sites did not increase the cytolytic activity of truncated HALTs-4 and they may play a role in the regulated secretory pathway.

## Author Contributions

W.Y.Y conducted all experiments and drafted the manuscript, J.S.H contributed to the design of work and revised manuscript.

